# Targeted memory reactivation during non-rapid eye movement sleep enhances neutral, but not negative, components of memory

**DOI:** 10.1101/2023.05.26.542120

**Authors:** Dan Denis, Jessica D. Payne

## Abstract

Emotionally salient components of memory are preferentially remembered at the expense of accompanying neutral information. This emotional memory trade-off is enhanced over time, and possibly sleep, though a process of memory consolidation. Sleep is believed to benefit memory through a process of reactivation during non-rapid eye movement sleep (NREM). Here, targeted memory reactivation (TMR) was used to manipulate the reactivation of negative and neutral memories during NREM sleep. Thirty-one male and female participants encoded scenes containing either a negative or neutral object superimposed on an always neutral background. During NREM sleep, sounds associated with these scenes were replayed, and memory for scene components was tested the following morning. We found that TMR during NREM sleep improved memory for neutral, but not negative scene objects. This effect was associated with sleep spindle activity, with a larger spindle response following TMR cues predicting TMR effectiveness for neutral items only. These findings therefore do not suggest a role of NREM memory reactivation in enhancing the emotional memory trade-off across a 12-hour period but do align with growing evidence of spindle-mediated memory reactivation in service of neutral declarative memory.

**Significance statement:** Memory reactivation during sleep is believed to be a key mechanism facilitating consolidation, the strengthening and stabilisation of memories over time. Emotional memories appear to be preferentially consolidated compared to neutral information, but the role sleep-related memory reactivation in this process is still unclear. Here, we found that experimentally reactivating memories during sleep in humans did not preferentially enhance emotional memory components but did improve neutral memory when tested after one night of sleep. These findings speak against a role of memory reactivation in the early stages of memory consolidation.

## Introduction

Intrinsically emotional experiences occupy a privileged place in our memory. Emotional events more strongly capture our attention (Yiend, 2010), are remembered more vividly (Kark and Kensinger, 2019), and are more resistant to forgetting compared to neutral information (Yonelinas and Ritchey, 2015). So strong is this effect that emotionally salient aspects of an event are preferentially remembered at *the expense* of accompanying neutral information (Kensinger et al., 2007). Numerous studies have documented this emotional memory trade-off effect, where negative aspects of scenes are preferentially remembered at the expense of accompanying neutral components, with no such effect being shown for completely neutral scenes (e.g. Denis et al., 2022; Kensinger et al., 2007; Payne et al., 2008; Sopp et al., 2018; Waring et al., 2010).

At initial encoding, an array of neurophysiological processes differentiate emotional from neutral stimuli (Phelps and LeDoux, 2005; Shields et al., 2019; Kim and Payne, 2020). Despite these immediate differences, the emotional memory bias becomes magnified over time, as memory performance diverges for emotional and neutral information after a delay of at least a few hours compared to immediate memory testing (Dolcos et al., 2005; Sharot and Yonelinas, 2008; Payne et al., 2012). This suggests that a period of memory consolidation, the process of memory stabilisation and strengthening, prioritises emotionally salient material (Stickgold and Walker, 2013; Kim et al., 2020; Cowan et al., 2021).

Sleep is an optimal state for memory consolidation to occur, as incoming sensory information is diminished (Brodt et al., 2023). A wealth of behavioural evidence shows enhanced memory retention over a period sleep compared to an equivalent amount of time spent awake (Berres and Erdfelder, 2021), even when the amount of wake-associated interference is equated (Talamini et al., 2008; Payne et al., 2012; Denis et al., 2020). Emerging evidence points to a role of memory reactivation during non-rapid eye movement sleep (NREM) underlying sleep’s beneficial effect on memory (Klinzing et al., 2019). During NREM, memories are repeatedly reactivated in hippocampal-neocortical circuits, a process clocked by the temporal coupling of hippocampal sharp-wave ripples (∼80-150Hz), thalamocortical sleep spindles (∼12-15Hz), and neocortical slow oscillations (∼1Hz) (Latchoumane et al., 2017; Cairney et al., 2018; Zhang et al., 2018; Kim et al., 2019; Schreiner et al., 2021).

A recent framework proposes that these mechanisms of NREM consolidation are biased towards emotional salient information (Cowan et al., 2021). In other words, memory reactivation, coordinated by SO-spindle coupling, should favour the reactivation of emotional memories. The emotional memory trade-off effect appears to be enhanced over a night of sleep compared to a day awake (Payne et al., 2008), an effect that has been frequently replicated (Payne and Kensinger, 2011; Payne et al., 2012, 2015; Cunningham et al., 2014; Alger et al., 2018), including in a recent large scale study of 280 participants (Denis et al., 2022, though see Bennion et al., 2015, 2017 for notable non-replications). These results occur against the backdrop of recent meta-analyses that have been unable to detect such effects when assessing the broader literature, collapsing across experimental designs and task types (Lipinska et al., 2019; Schäfer et al., 2020). It may be the case that the trade-off effect within individual items is more sensitive to the effects of sleep than other commonly used tasks. (Davidson et al., 2021), making it the ideal task to explore the role of reactivation processes in emotional memory consolidation.

Across the existing literature, no consistent correlations between any aspect of sleep and emotional memory have been found (Davidson et al., 2021). With regards to the emotional memory-trade off, some studies have found NREM sleep and sleep spindles to correlate with the magnitude of the trade-off, speaking in favour of NREM reactivation processes (Payne et al., 2015; Alger et al., 2018). On the other hand, a different study found correlations with rapid eye movement sleep (REM) (Payne et al., 2012), in line with other lines of research suggesting that REM sleep, and REM theta (4-8Hz) oscillations in particular, selectively enhance emotional memories (Nishida et al., 2009; Sopp et al., 2017; Kim et al., 2020).

Such studies are correlational in nature, and do not directly test the process of reactivation. Targeted memory reactivation (TMR) is an experimental technique through which the reactivation of certain memories can be directly manipulated. In this paradigm, unique sounds are paired with stimuli during encoding, with half being replayed during subsequent sleep. These sounds induce increases in sleep spindle activity, and the reinstatement of memory-related content (Bendor and Wilson, 2012; Rothschild et al., 2017; Belal et al., 2018; Cairney et al., 2018; Schreiner et al., 2018; Wang et al., 2019; Ngo and Staresina, 2022). Meta-analytic evidence has shown that memory for items that were reactivated during sleep is enhanced compared to the un-reactivated memories (Hu et al., 2020). The application of TMR to the question of emotional memory has yielded mixed results. While some studies suggest that emotional memories do benefit from reactivation in NREM (Cairney et al., 2014; Lehmann et al., 2016; Yuksel et al., 2023), null findings have also been reported (Rihm and Rasch, 2015; Ashton et al., 2018; Hutchison et al., 2021; Pereira et al., 2022). Similar mixed results have been reported when TMR is applied during REM (Sterpenich et al., 2014; Lehmann et al., 2016; Hutchison et al., 2021; Yuksel et al., 2023). However, no studies to date have examined the effect of TMR on the emotional memory trade-off effect (i.e., does TMR enhance emotional information at the expense of accompanying neutral information?).

The present study set out to test the hypothesis that reactivation during NREM sleep facilitates the selective enhancement of negative aspects of memory. By combining the emotional memory trade-off task, which may be more sensitive to sleep effects compared with other tasks(Davidson et al., 2021), and TMR to manipulate memory reactivation, the following specific hypotheses were tested: 1) Negative components of memory would be preferentially enhanced at the expense of accompanying neutral components; 2) This emotional memory trade-off effect would be further enhanced by TMR during NREM sleep; 3) The effect of TMR would be associated with TMR-evoked sleep spindle activity. As an exploratory aim, correlational analyses between ongoing oscillatory activity during sleep with emotional memory were performed. These analyses focused on SO-spindle coupling during NREM sleep, based on prior work suggesting oscillatory coupling to be a mechanistic driver of consolidation (Klinzing et al., 2019), and REM theta oscillations, as a putative marker of REM-based emotional memory consolidation (Hutchison and Rathore, 2015).

## Methods

### Participants

Participants were 31 undergraduate students recruited at the University of Notre Dame (M_age_ = 20 years, SD_age_ = 1 year; 68% female). See **Table S1** for sample demographics. Inclusion criteria for the study included no self-reported history of any sleep, neurologic, or psychiatric disorder, and a self-reported bedtime of no later than 2am and sleeping on average for no less than six hours per night. All participants were instructed to keep a regular sleep schedule for the three nights prior to the study, and to abstain from caffeine the day of the study. All participants provided informed consent prior to taking part in the study and were compensated with either cash or course credit. Recruitment was through advertisement of the study on SONA and flyers placed around the University of Notre Dame campus. The study received ethical approval from the University of Notre Dame Institutional Review Board.

### Design

#### Study overview

The study design is depicted in **Figure 1**. After providing informed consent, participants filled out questionnaires regarding their subjective sleep habits over the past three nights, general subjective sleep quality over the past month as assessed by the Pittsburgh Sleep Quality Index (Buysse et al., 1989), and their diurnal preference as assessed by the Morningness-Eveningness Questionnaire (Horne and Ostberg, 1976). Following this, participants were wired for EEG (see below). They then reported their subjective levels of alertness via the Stanford Sleepiness Scale (Hoddes et al., 1972), before completing the incidental encoding portion of the emotional memory trade-off task. At approximately 11pm, participants went to bed and had a nine-hour sleep opportunity, before being awoken at 8am the next day. During the first hour of NREM (N2+N3) sleep, half of the sounds were replayed (see Targeted memory reactivation below). In the morning, participants had the EEG cap removed and were given the opportunity to shower. They then completed a second Stanford sleepiness scale assessment, before completing the recognition portion of the emotional memory trade-off task. See **Table S1** for all subjective sleep and alertness measures.

**Figure 1.**
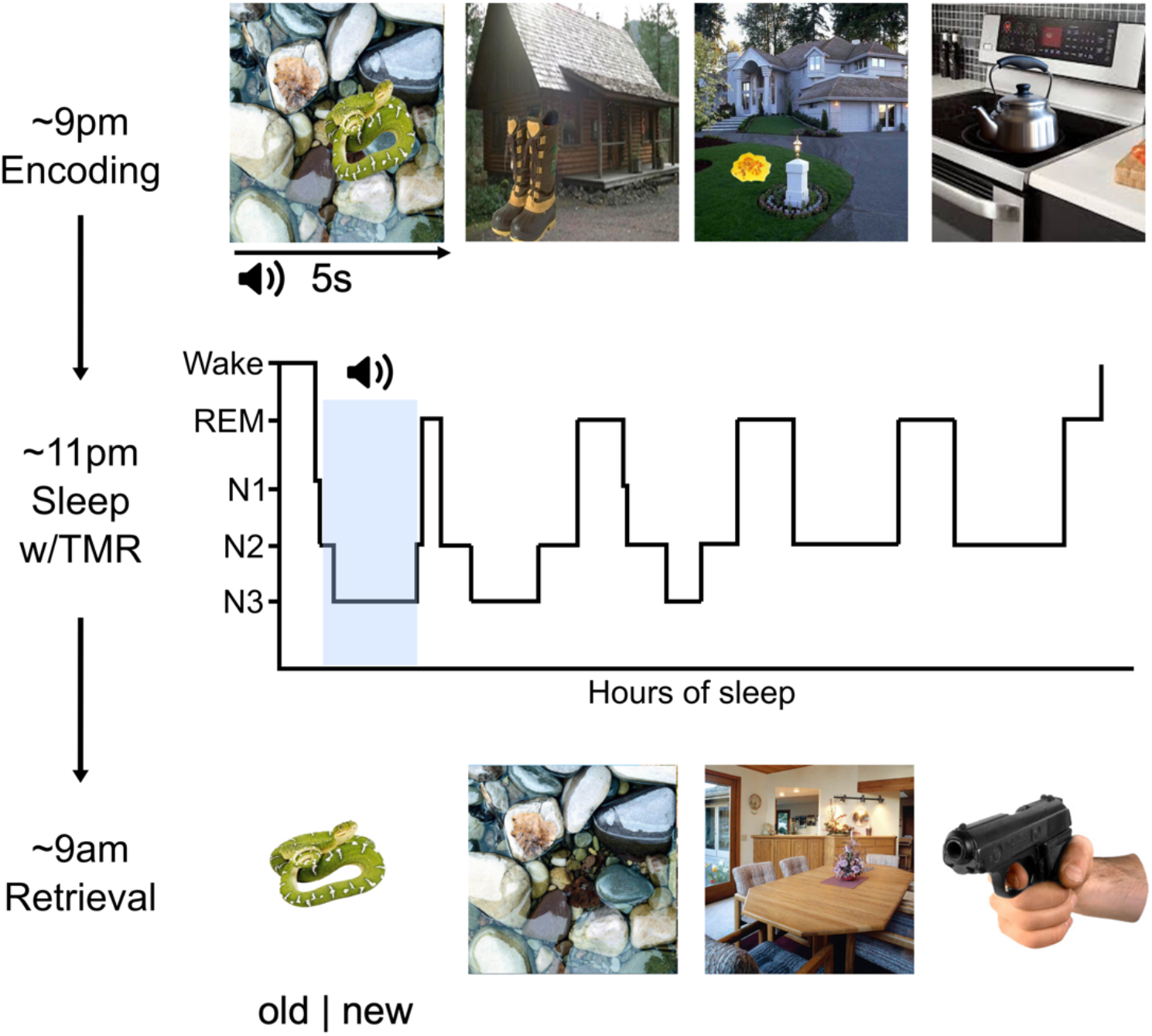
Experimental design. In the evening, participants incidentally encoded 92 scenes containing either a negative or neutral object superimposed on an always neutral background. Each scene was accompanied by a sound naturally linked to the object. During the first hour of non-rapid eye movement sleep, half of the sounds were replayed. A surprise memory test was administered the following morning. Participants viewed scene components individually and had to indicate whether the component was old or new.

#### Emotional memory trade-off task

The studied materials consisted of 92 scenes depicting negative (n = 46) or neutral (n = 46) objects placed on plausible, always neutral backgrounds. Each scene was accompanied by a sound that was conceptually related to the object component of the scene (e.g. a “woof” sound was paired with a scene depicting a dog in a park). The trade-off scenes used were taken from a larger database of scenes and have been used in previous studies (Denis et al., 2022; Cunningham et al., 2021). The sounds were taken from a prior unpublished report (Personal communication, 2019) and from the website Freesound (https://freesound.org/).

The emotional memory trade-off task consists of an initial incidental encoding portion followed by a surprise recognition test after the delay (**Figure 1**). During encoding, participants viewed each of the 92 scenes. Each trial began with a fixation cross displayed in the centre of the screen for 1s (+/- 200ms inter-trial jitter) before the scene was displayed on the screen for 5s. The associated sound was played at the onset of each trial. Participants were instructed to rate the scene for its perceived valence and arousal. At the end of each trial, participants rated each scene on a 1-5 scale for its valence (very negative to very positive) and arousal (not at all arousing to very arousing). After a 2s inter-trial interval, the next trial began. Participants were instructed to remain still and relaxed, and to limit eye blinks and movements while the scene was on the screen. There was a break halfway through to allow participants to stretch and adjust their posture. At no point during the encoding session were participants informed that their memory would be tested.

Following the sleep period, participants completed a surprise, self-paced recognition test in which the objects and backgrounds were presented separately and one at a time. The old objects and backgrounds that comprised the full scenes at encoding were intermixed with an equal number of new object and background components of scenes that were not viewed at encoding. For each trial, participants were instructed to indicate whether the component was old or new as quickly and as accurately as possible using the keyboard. There was a total of 368 trials at recognition (92 *old* objects (46 negative, 46 neutral), 92 *old* backgrounds (46 originally paired with negative objects, 46 originally paired with neutral objects), 92 *new* objects (46 negative, 46 neutral), and 92 *new* backgrounds (all neutral). No sounds were played at the recognition test.

#### Targeted memory reactivation

Soft pink noise was played out of a bedside speaker throughout the sleep period. When participants reached five-minutes of uninterrupted N2 or N3 sleep, cueing ensued. A total of 46 randomly selected sounds (23 negative, 23 neutral) that had been presented with scenes during encoding were played, along with six (3 negative, 3 neutral) controls sounds that were not associated with any of the scenes presented at encoding. All 52 sounds were unobtrusively presented in a random order, with a five second inter-stimulus interval. After all 52 sounds were presented, the order was re-randomised and a new cueing block began. Cues were presented during both N2 and N3 sleep, and continued for one hour or until the first period of rapid eye movement sleep was reached. Cueing immediately stopped upon sign of an arousal or shift to N1 sleep or wakefulness. In these cases, cueing resumed after re-entering N2 or N3 sleep.

### EEG

#### Acquisition

EEG was collected from all participants during both the encoding and sleep portions of the experiment at a sampling rate of 500Hz. For this analysis, we focused on just the sleep EEG recordings. Data were acquired from 58 electrodes positioned according to the 10-20 system. Additional electrodes were placed on the left and right mastoid (for offline re-referencing), above the right and below the left eye (for EOG measurements), and two placed on the chin (for EMG measurements). A BrainVision actiChamp amplifier and Recorder software was used to acquire the data. Impedances were kept below 15 kΩM. EEG data were downsampled to 250Hz for all offline analyses. Data were sleep scored offline according to standard criteria (Iber et al., 2007). Sleep architecture is shown in **Table S2**.

#### EEG Preprocessing

Data were downsampled to 250Hz. Bad channels were identified by eye and interpolated using a spherical splines algorithm. Next, EEG data were notch filtered at 60Hz, high pass filtered at 0.3Hz, and re-referenced to the average of the two mastoid channels. Data were then epoched from -1 to 3 seconds around the onset of all cues that were presented during either N2 or N3 sleep. Bad epochs were identified as those exceeding a voltage threshold of +/- 500 uV, and using the joint probability function in EEGLAB, with a threshold of six standard deviations for single channels and two standard deviations for the global signal (all channels grouped). Finally, each record was visually inspected to ensure adequacy of the artefact rejection procedure.

#### TMR analysis

Clean, artefact free epochs were baseline adjusted to the mean voltage in the 1 second before cue onset. Complex Morlet wavelets were then used to decompose the time series data into time-frequency representations (TFRs). Spectral power was extracted at 30 logarithmically spaced frequencies from 2-40 Hz with the number of wavelet cycles increasing from 5 to 10 in 30 logarithmically spaced steps to match the number of frequency bins. Time frequency power was averaged over trials, and decibel normalised within-participant, where the baseline was mean power in the 500-200 ms prior to cue-onset. This choice of baseline was chosen so as to mitigate contamination of the baseline period by post-stimulus activity. As in prior TMR experiments, analyses were performed at electrode Cz.

Sleep spindles that occurred following TMR cues were detected using an automated algorithm (see below for details). For each spindle that was detected following a TMR cue, its amplitude was extracted. To calculate the probability of a spindle occurring following a TMR cue, established procedures were followed (Schechtman et al., 2021; Liu et al., 2023). For each trial, a spindle value of 1 was assigned to the time points where a spindle was detected, and 0 where no spindle was detected. The spindle probability was then computed as the mean spindle values across trials for each time point.

#### Slow oscillation-spindle coupling

SO-spindle coupling was detected at all EEG electrodes during artifact-free NREM sleep using well-validated approaches. First, individual channels where are least one Hjorth parameter was > 3 times the standard deviation of the mean for at least 25% of all epochs was considered bad and was interpolated. Second, individual epochs were removed if > 50% of channels had at least one Hjorth parameter > 3 times the standard deviation of the mean. For epochs where < 50% of channels were above the threshold, these channels were interpolated on an epoch-by-epoch basis. For both steps, artefact detection was performed over two iterations.

Sleep spindles were detected at all electrodes using a wavelet based detector (Wamsley et al., 2012; Warby et al., 2014). As a first step, we identified each participant’s fast spindle peak frequency from their NREM power spectrum. Power spectral density (PSD) was estimated using Welch’s method with 5s Hamming windows and 50% overlap. PSD estimates were obtained from the derivative of the EEG time series to minimise 1/*f* scaling and maximise spectral peaks (Cox et al., 2017). The largest, most prominent peak within a broadly defined fast spindle range of 12.5 – 16Hz was taken as that individual’s peak spindle frequency. After detecting each individual’s spindle peak frequency, the EEG signal was subject to a time-frequency decomposition using complex Morlet wavelets. The wavelet parameters were tuned for each individual based on their spindle peak. Specifically, the peak frequency of the wavelet was set at that individual’s spindle peak, with a 3Hz bandwidth (full-width, half-max) centred on the peak frequency (Denis et al., 2021b). Spindles were detected by applying a thresholding algorithm to the extracted wavelet scale. A spindle was detected whenever the wavelet signal exceeded a threshold of nine times the median signal amplitude of all artefact-free data for a minimum of 400ms. The threshold of nine times the median empirically maximises between class (“spindle” vs “non-spindle”) variance in previous samples of healthy participants with 12-15 Hz overnight spindles (Mylonas et al., 2020).

Slow oscillations were detected using a second automated detector (Staresina et al., 2015). First, data were bandpass filtered between 0.5 - 4Hz and all positive-to-negative zero crossings were identified. Candidate slow oscillations were marked if two such consecutive zero crossings fell 0.8 - 2 seconds apart, corresponding to 0.5 - 1.25 Hz. Peak-to-peak amplitudes for all candidate oscillations were determined, and oscillations in the top quartile (i.e., with the highest amplitudes) at each electrode were retained as slow oscillations (Staresina et al., 2015; Helfrich et al., 2018).

To identify SO-spindle coupling events, EEG data were bandpass filtered in individualised spindle frequency range. Then, the Hilbert transform was applied to extract the instantaneous phase of the delta (0.5 - 4 Hz) filtered signal and the instantaneous amplitude of the spindle filtered signal. For each detected spindle, the peak amplitude of that spindle was determined. It was then determined whether the spindle peak occurred within the time course (i.e., between two positive-to-negative zero crossings) of any detected slow oscillation. If the spindle peak was found to occur during a slow oscillation, the phase angle of the slow oscillation at the peak of the spindle was calculated. We extracted the density of coupled spindles (calculated as the number of slow oscillation-coupled spindles per minute of NREM sleep) and the average coupling phase (in degrees) and consistency (vector length).

#### REM power spectral density

To estimate REM theta power spectral density (PSD), artefactual REM epochs were removed using the same automated artefact rejection procedure as above. REM PSD estimates were obtained at each EEG channel using the same method as for spindle peak detection. Given that signal amplitude is at least partly driven by individual difference factors such as skull thickness and gyral folding (Cox and Fell, 2020), we then normalised, within participant, each electrode’s power spectrum by dividing power at each frequency by that electrode’s average power (Denis et al., 2021a). Each individual’s peak frequency within a broadly defined theta range (3-8Hz) was identified from the power spectrum. Theta PSD estimates were then calculated by averaging together PSD in frequencies ranging +/- 1.5Hz around the theta peak (Harrington et al., 2020).

### Memory analysis

To assess memory performance, we calculated a corrected recognition score in line with previous research utilising the emotional memory trade-off task (e.g. Cunningham et al., 2021; Denis et al., 2022). For this measure, a “hit” was defined as saying “old” to an old trial, and a false alarm was defined as saying “old” to a new trial. For each scene component, we calculated the proportion of hits and false alarms and then subtracted the false alarm rate from the hit rate to obtain the corrected recognition score. For the purposes of correlating TMR cue-evoked EEG activity with performance, we also calculated the cueing benefit by subtracting corrected recognition scores for the uncued items from corrected recognition scores for the cued items (Groch et al., 2017). As such, a more positive value reflects a greater efficacy of TMR in terms of improving memory.

### Statistical analysis

#### Behaviour

The effect of TMR on the emotional memory trade-off was assessed via a 2 (emotion: negative, neutral) x 2 (component: object, background) x 2 (TMR: cued, uncued) linear mixed effects model with participant entered as a random effect, and corrected recognition scores as the dependent variable. Follow-up estimated marginal means tests were used as appropriate.

#### EEG

We first contrasted time-frequency responses to memory and control cues by performing paired t-tests across all time and frequency points. To ensure that any differences between conditions were not due to differences in trial counts, the number of trials was matched between conditions by randomly selecting a subset of trials from the higher trial count condition. A similar approach was used to contrast negative and neutral cues. To control for multiple comparisons across time and frequency points we employed a cluster-based permutation approach (Maris and Oostenveld, 2007). For each analysis, clusters were formed from adjacent time-frequency points that met an uncorrected threshold of *p* < .05. Permutation distributions were then created by randomly shuffling the condition labels 1000 times and retaining the cluster with the maximum statistic for each permutation. A cluster-corrected *p* < .05 was deemed statistically significant.

To assess associations between cue-evoked EEG activity and the TMR benefit, time-frequency points in significant clusters were averaged together to obtain a single value for each cluster reflecting memory/emotion-sensitive activity. For each cluster separately, we regressed cueing benefit against cluster power, emotion (Negative, Neutral), and their interaction. Robust regression procedures were used to minimise the influence of outliers.

To examine relationships between endogenous slow oscillation-spindle coupling/REM theta PSD and memory, we also used robust linear regressions. In separate models, we regressed corrected recognition scores against SO-spindle coupling/REM theta PSD at each electrode, with emotion (Negative, Neutral) and the interaction term included as fixed effects. Multiple comparisons across electrodes were controlled for using the same cluster-based permutation testing framework described above. Circular-linear correlations between spindle coupling phase and corrected recognition scores were run at each electrode using the *circStat* toolbox (Berens, 2009), with multiple comparisons again being controlled for via cluster-based permutation tests. For coupling phase analysis, negative and neutral memory was analysed separately.

## Results

### Behaviour

At encoding, participants rated negative scenes as significantly more negatively valanced and significantly more emotionally arousing compared to neutral scenes (valence: negative M = 2.57, SD = 0.41, neutral M = 4.45, SD = 0.40; arousal: negative M = 4.49, SD = 1.04, neutral M = 3.40, SD = 0.83; all *p*s < .001). Subjective valence and arousal ratings did not correlate with any aspect of subsequent memory (all *p* > .20).

We next examined memory scores at the post-sleep recognition test (see **Table S3** for raw memory scores). A significant interaction between emotion and component (*F* (1, 210) = 72.86, *p* < .001) indicated the presence of an emotional memory trade-off effect (**Figure 2A**). As expected, memory for negative objects (M = .75, SD = .16) was significantly better than memory for neutral objects (M = .64, SD = .20); *t* (210) = 6.07, *p* < .001. This coincided with significantly *worse* memory for backgrounds initially paired with a negative object (M = .49, SD = .19) compared to backgrounds initially paired with a neutral object (M = .60, SD = .19); *t* (210) = -6.00, *p* < .001.

**Figure 2.**
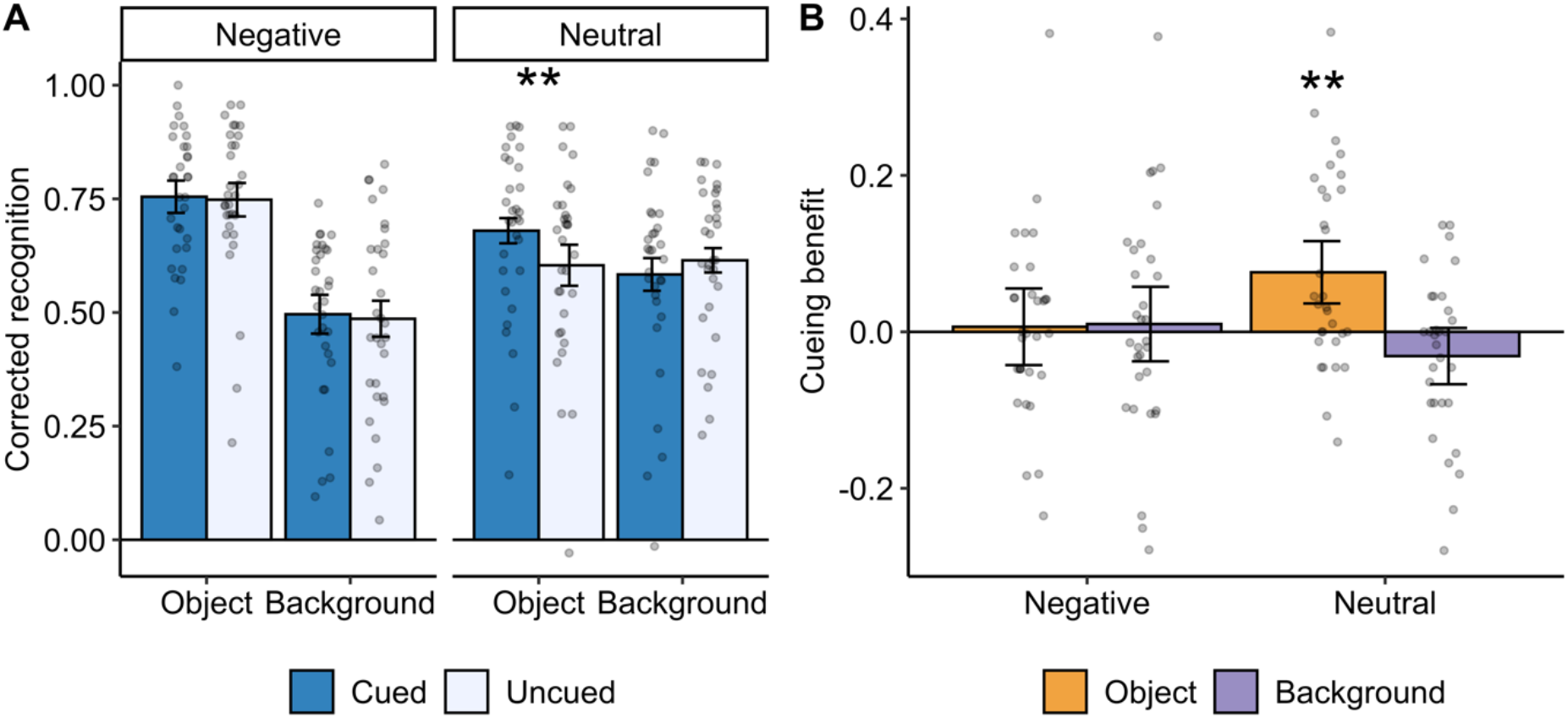
Behavioural results. **A** – Component corrected recognition scores for cued (dark blue) and uncued (light blue) items. **B** – Cueing benefit (cued minus uncued corrected recognition). All error bars indicate 95% confidence intervals. ** = *p* < .01.

The overall emotional memory trade-off was superseded by a significant 3-way interaction between TMR, emotion, and component (*F* (1, 210) = 4.71, *p* = .03), suggesting a selective effect of TMR on memory (**Figure 2A**). Contrary to our hypothesis, there was no difference in memory between negative objects that were either cued (M = .76, SD = .15) or uncued (M = .75, SD = .17); *t* (210) = 0.25, *p* = .80. On the other hand, memory for neutral objects that were cued during sleep (M = .68, SD = .19) was significantly enhanced relative to uncued neutral objects (M = .60, SD = .21); *t* (210) = 2.99, *p* = .003. TMR did not alter memory for backgrounds paired with either negative or neutral objects (*p*s > .61). We note these results remained significant after removal of a potential outlier (neutral memory > 1.5 * the inter-quartile range).

The TMR benefit for neutral objects was significantly greater than zero (*t* (30) = 3.40, *p* = .002), with TMR facilitating a 7.6% improvement in memory for neutral objects (**Figure 2B**). At the individual level, 19 (61%) of 31 participants exhibited a beneficial effect (i.e., > 0) of TMR on neutral object memory. Although TMR significantly improved neutral object memory, memory for cued neutral objects was still significantly worse than memory for negative objects (*t* (210) = 2.93, *p* = .004).

It is possible that, rather than enhancing memory for cued items, TMR suppresses or impairs memory for the uncued items. To test for this possibility, we compared memory for the uncued items against a separate sample who underwent the exact same procedure, except no TMR cues were delivered during sleep (**Figure S1**). There was no main effect of group, nor any interactions involving group (all *p*s > .15). As such, uncued memories showed similar to performance to what occurs over a night of normal sleep with no external cueing.

### Cue-evoked EEG activity during sleep

We next turned our attention to EEG activity following cue presentation during sleep (number of cues by condition and sleep stage presented in **Table S4**). We first tested for memory-specific processing by contrasting TFRs to sounds originally paired with a scene during encoding (i.e., memory related cues) to non-learning control sounds. Memory-related cues evoked significantly larger responses in two time-frequency clusters (**Figure 3A**). The first cluster emerged 358-1016ms following cue onset in canonical theta-band frequencies ranging from 4.57-7.65Hz (*t*_sum_ = 3445, *p* = .026). A second cluster formed between 812-1282ms at frequencies broadly corresponding to the sleep spindle band (14.24 – 19.41Hz; *t*_sum_ = 1923, *p* = .041). As such, the brain response to sounds cued during sleep was sensitive to whether that sound was associated with a scene viewed during pre-sleep memory encoding.

**Figure 3.**
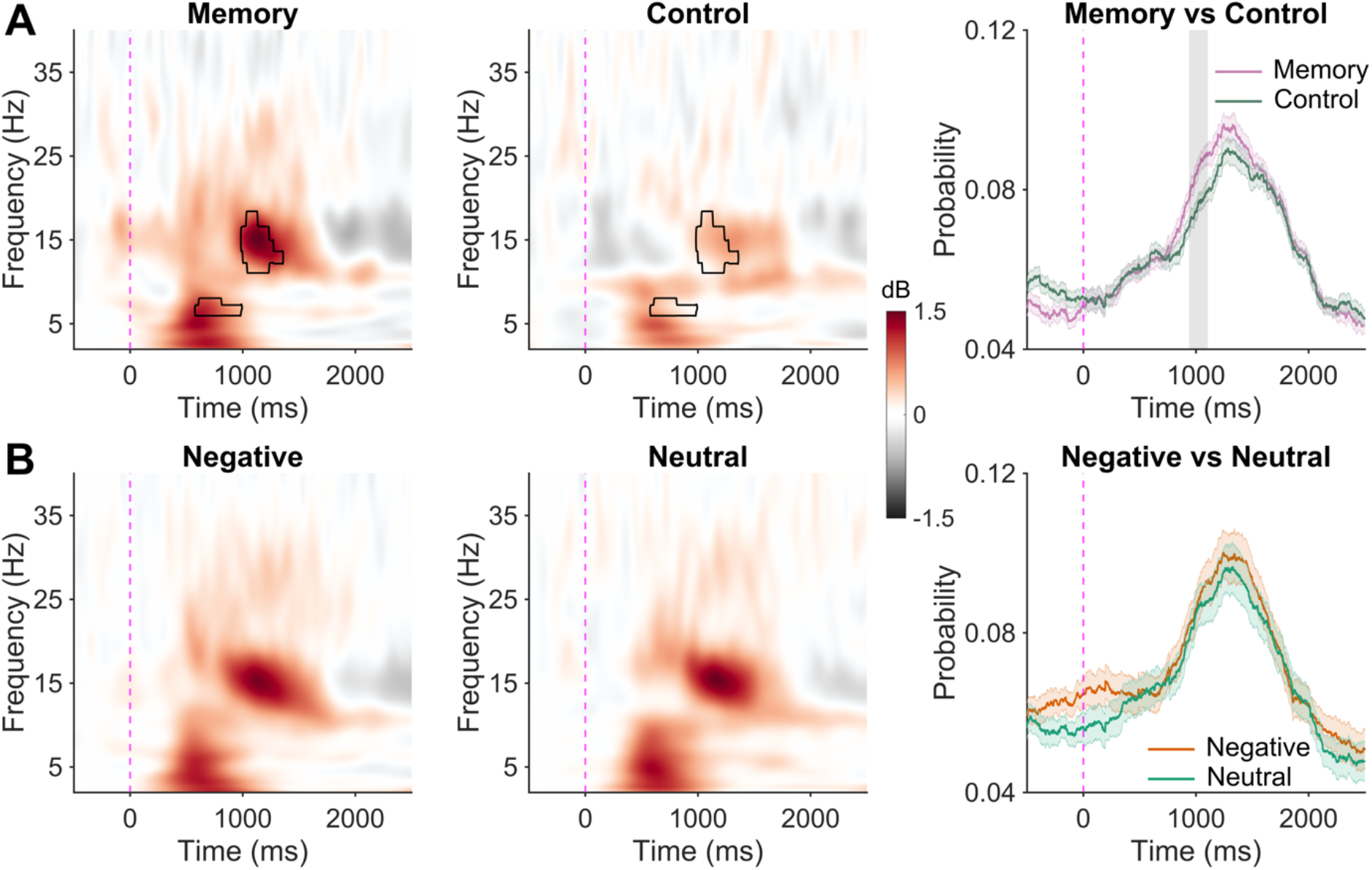
Cue-elicited EEG activity during sleep. **A** - EEG response to memory and control cues. Time 0 indicates cue onset. Left: Time-frequency representation (TFR) of cue-elicited neural activity following memory-related cues. Middle: TFR of response to control sounds. Significant differences between memory and control cues (cluster corrected) highlighted in black. Right: Spindle probability following either memory (pink) or control (green) cues. Significant difference (cluster corrected) highlighted in grey. Shaded area depicts the 95% confidence interval. **B** - Same as A, but for either negative (left) or neutral (middle) cues. Right hand plot shows spindle probability following either negative (orange) or neutral (green) cues. Shaded area depicts the 95% confidence interval.

The enhanced power in spindle frequencies could either reflect an increased occurrence of spindles following memory related cues, higher spindle amplitude following memory cues, or both. Regarding occurrence, there was a significantly higher probability of a spindle occurring following a memory-related cue compared to a control cue 876-1208ms following sound onset (*t*_sum_ = 605, *p* = .008), a time-window closely corresponding to cue-evoked increase in spindle power (**Figure 3A**). On the other hand, there was no difference in spindle amplitude following memory (M = 58.24µV, SD = 9.86µV) or control (M = 58.48µV, SD = 9.81µV) cues (*t* (58) = 0.10, *p* = .92).

Having established brain responses specific to memory-related TMR cues, we next examined TFR responses to memory-related negative cues and memory related neutral cues (**Figure 3B**). No significant clusters emerged. Similarly, there were no significant differences in either spindle probability or amplitude. As such, it appears that the valence of the cued memory did not impact on the EEG dynamics following sound presentation.

To test our third hypothesis, we correlated cluster-averaged power in the spindle-band cluster with the benefit of TMR. There was a significant interaction between spindle cluster power and emotion on the TMR benefit (*F* (52) = 5.51, *p* = .023; **Figure 4**). Cue-evoked spindle power correlated positively with the cueing benefit for neutral objects (*r* = .54, *p* = .004) but not negative objects (*r* = .02, *p* = .58). There was no main effect of theta cluster power on the TMR benefit (*F* (1, 52) = 0.001, *p* = .97), nor was there an interaction with emotion (*F* (1, 52) = 0.02, *p* = .89).

**Figure 4.**
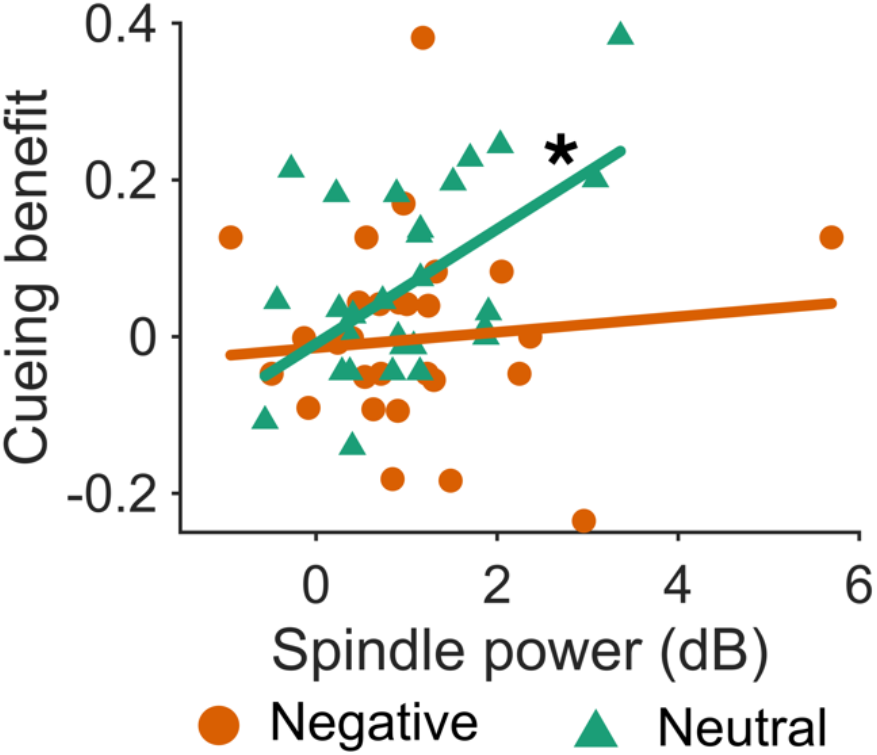
Cue-evoked spindle power is associated with the cueing benefit of neutral, but not negative, object memory. * = *p* < .05.

### Endogenous sleep physiology correlations

To examine our exploratory aim, we performed correlational analyses between ongoing, endogenous oscillations believed to facilitate consolidation, and object memory. For all measures assessed (spindle density, coupled spindle density, spindle coupling phase and consistency, REM theta power), no main effects or interactions with emotion on memory were found (*p*s > .11). Additionally, no main effects or interactions was observed for N3 time, NREM time (N2 + N3), or REM time (*p*s > .12).

## Discussion

This study tested the role of memory reactivation during NREM sleep on enhancing negative components of memory. Although we observed a substantial emotional memory trade-off, memory reactivation did not enhance this trade-off further. Instead, reactivation improved memory for neutral objects, an effect mediated by sleep spindle activity. These results further support findings that neutral declarative memories are reactivated and consolidated during NREM sleep, with these consolidation effects being mediated by spindle effects. We extend the existing literature by showing that these same processes do not appear to enhance emotional memories across a twelve-hour delay. We consider several potential reasons as to why TMR did not enhance the emotional memory trade-off effect.

First, it is possible that overnight enhancement of the trade-off is REM-dependent, which would be in line with a prior report utilising the same task in an overnight design (Payne et al., 2012) and other work implicating REM theta spectral power in selective emotional memory processing (Hutchison and Rathore, 2015). We ruled out an effect of REM sleep in the present study, with no correlations between either REM time or REM theta power being found with emotional memory. This null effect agrees with the larger literature that has not found consistent links between REM sleep and emotional memory (Davidson et al., 2021).

A second is explanation could be that because emotional memories were prioritised for consolidation (Kim and Payne, 2020; Cowan et al., 2021), these memories were being reactivated endogenously, with no additive effect of additional reactivations via TMR. This could explain why neutral memories were benefitted by TMR, as these memories would not have been reactivated as often endogenously. Memory reactivation during NREM sleep is closely tied with SO-spindle coupling events (Schreiner et al., 2021), therefore under this explanation it would be expected that ongoing SO-spindle coupling would positively correlate with emotional memory. However, no such associations were seen in the present study.

Memory consolidation is a slow process that continues in the weeks and months following encoding (Dudai, 2012). Sleep’s impact on emotional memory can be seen years after the original experience (Wagner et al., 2006). Given how powerful the effect of emotion is, it is possible that the benefits of memory reactivation on emotional memory might not be visible after the 12-hour delay employed in this study. Future studies should seek to examine whether memory reactivation makes emotional memories more resistant to forgetting over longer intervals. A related explanation is that participants were at a functional ceiling with regards to negative object memory, limiting the capacity to which TMR could further enhance these memories. As well as increasing the length of delay, a higher number of items during encoding could also serve to weaken memory, which may heighten the observed TMR benefits (Creery et al., 2015; Cairney et al., 2016).

One implication of our findings is that TMR can be used to enhance otherwise de-prioritised memories (i.e., neutral relative to negative memories). A similar finding was reported by Oudiette and colleagues who found that TMR improved memory for low reward memories, but a natural bias towards high reward memories was not boosted by TMR (Oudiette et al., 2013). Together, these findings show that TMR is able to increase the capacity of sleep-associated memory consolidation, presumably through eliciting memory reactivations that would not have occurred endogenously. An intriguing clinical application would be to investigate whether TMR can be used to boost memories that are maladaptively de-prioritised, such as positive information in major depression (Talamini and Juan, 2020; Everaert et al., 2022).

Some limitations of the work should be considered. Although we did not find any correlations with REM sleep, we did not include a REM TMR group. We focused on NREM reactivation here for three reasons: 1) A large evidence base already exists for an effect of NREM reactivation on memory consolidation in general (Klinzing et al., 2019); 2) Recent models of adaptive memory processing during sleep have placed a clear emphasis on NREM sleep processes (Cowan et al., 2021), and 3) Prior TMR studies of emotional memory have found no consistent benefit of reactivating during REM, with one study even finding it induces forgetting of emotional memories (Yuksel et al., 2023). A second limitation is that the relatively low number of trials and high levels of performance for negative items precluded analyses of subsequently remembered vs subsequently forgotten trials, which may have revealed further insights into how memories were being re-processed following TMR.

Taken together, the current study provides further evidence that memory reactivation during sleep improves memory for neutral declarative information, an effect mediated by sleep spindle activity. This process does not appear to support the same function for emotional memories over the first 12 hours following encoding. This study is the first to attempt to enhance the emotional memory trade-off via a sleep manipulation and future research should now seek to better understand how memory reactivation during sleep may support this trade-off over longer time scales.

## Acknowledgements

This work was supported by the National Science Foundation under Grant BCS-2001025, awarded to J.D.P. We would like to thank James W. Antony for providing us with some of the sound clips used in the experiment.

---

**Table S1.** Demographics and subjective sleep and alertness measures

|  | M | SD |
| --- | --- | --- |
| Age (years) | 20 | 1.3 |
| Sex |  |  |
| Female (%) | 68 |  |
| Male (%) | 32 |  |
| PSQI | 4.5 | 2.80 |
| MEQ | 45.7 | 8.56 |
| Sleep log bedtime | 01:22 | 01:27 |
| Sleep log rise time | 09:03 | 01:22 |
| Sleep log TST (mins) | 462 | 74 |
| Encoding subjective alertness | 2.47 | 0.98 |
| Recognition subjective alertness | 2.89 | 1.20 |

*Note.* M = Mean, SD = Standard deviation. Sex expressed as a percentage of participants. PSQI = Pittsburgh sleep Quality Index. Higher score indicates worse subjective sleep quality (theoretical range: 0-21). MEQ = Morningness-eveningness Questionnaire. Higher score indicates greater evening preference (theoretical range: 16-86). TST = Total sleep time. Sleep log measures are the average of the three nights prior to the experimental night.

**Table S2.** Sleep architecture on the experimental night

|  | M | SD |
| --- | --- | --- |
| Total sleep time (min) | 503 | 32 |
| Sleep onset latency (min) | 24.5 | 23.16 |
| Wake after sleep onset (min) | 33.91 | 15.86 |
| Sleep efficiency (%) | 90.62 | 4.85 |
| N1 time (% of TST) | 4.40 | 4.64 |
| N2 time (% of TST) | 54.60 | 7.68 |
| N3 time (% of TST) | 18.50 | 6.03 |
| REM time (% of TST) | 22.51 | 6.79 |

*Note.* M = Mean, SD = Standard deviation. TST = Total sleep time.

**Table S3.** Memory scores

|  | Cued |  | Uncued |  |
| --- | --- | --- | --- | --- |
|  | M | SD | M | SD |
| <u>Hit rate</u> |  |  |  |  |
| Negative objects | .88 | .09 | .87 | .12 |
| Negative backgrounds | .59 | .17 | .58 | .20 |
| Neutral objects | .77 | .15 | .69 | .16 |
| Neutral backgrounds | .67 | .19 | .71 | .15 |
| <u>False alarm rate</u> |  |  |  |  |
| Negative objects | .12 | .09 |  |  |
| Neutral objects | .09 | .08 |  |  |
| Backgrounds | .09 | .08 |  |  |

*Note.* M = Mean, SD = standard deviation.

**Table S4.** Number of sounds presented during sleep

|  | N2 |  | N3 |  | N2 + N3 |  |
| --- | --- | --- | --- | --- | --- | --- |
|  | M | SD | M | SD | M | SD |
| Negative | 81.31 | 64.03 | 165.97 | 70.37 | 247.28 | 52.96 |
| Neutral | 80.00 | 63.85 | 167.14 | 70.64 | 247.14 | 52.92 |
| Control | 35.55 | 28.83 | 76.52 | 32.59 | 112.07 | 24.72 |

*Note.* M = Mean, SD = standard deviation

**Figure S1.**
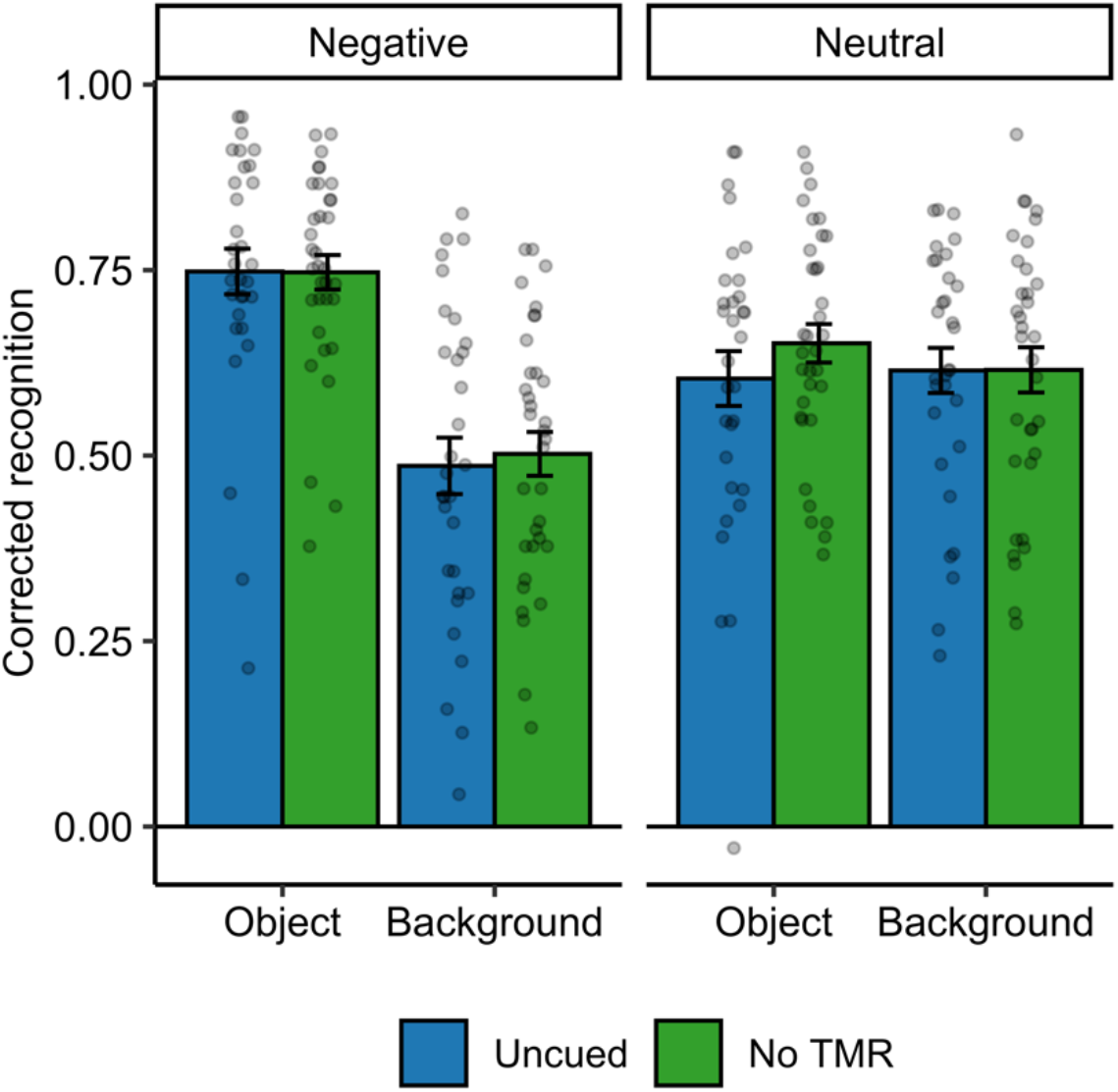
Component recognition scores for uncued items in the TMR (blue bars) compared with scores for a no TMR control group (green bars). Error bars indicate the between-subject standard error.

